# High-resolution awake mouse fMRI at 14 Tesla

**DOI:** 10.1101/2023.12.08.570803

**Authors:** David Hike, Xiaochen Liu, Zeping Xie, Bei Zhang, Sangcheon Choi, Xiaoqing Alice Zhou, Andy Liu, Alyssa Murstein, Yuanyuan Jiang, Anna Devor, Xin Yu

## Abstract

High-resolution awake mouse fMRI remains challenging despite extensive efforts to address motion-induced artifacts and stress. This study introduces an implantable radiofrequency (RF) surface coil design that minimizes image distortion caused by the air/tissue interface of mouse brains while simultaneously serving as a headpost for fixation during scanning. Furthermore, this study provides a thorough acclimation method used to accustom animals to the MRI environment minimizing motion induced artifacts. Using a 14T scanner, high-resolution fMRI enabled brain-wide functional mapping of visual and vibrissa stimulation at 100x100x200µm resolution with a 2s per frame sampling rate. Besides activated ascending visual and vibrissa pathways, robust BOLD responses were detected in the anterior cingulate cortex upon visual stimulation and spread through the ventral retrosplenial area (VRA) with vibrissa air-puff stimulation, demonstrating higher-order sensory processing in association cortices of awake mice. In particular, the rapid hemodynamic responses in VRA upon vibrissa stimulation showed a strong correlation with the hippocampus, thalamus, and prefrontal cortical areas. Cross-correlation analysis with designated VRA responses revealed early positive BOLD signals at the contralateral barrel cortex (BC) occurring 2 seconds prior to the air-puff in awake mice with repetitive stimulation, which was not detected using a randomized stimulation paradigm. This early BC activation indicated a learned anticipation through the vibrissa system and association cortices in awake mice under continuous training of repetitive air-puff stimulation. This work establishes a high-resolution awake mouse fMRI platform, enabling brain-wide functional mapping of sensory signal processing in higher association cortical areas.

**Significance Statement:** This awake mouse fMRI platform was developed by implementing an advanced implantable radiofrequency (RF) coil scheme, which simultaneously served as a headpost to secure the mouse head during scanning. A thorough acclimation method was used to accustom animals to the MRI environment minimizing motion induced artifacts. The ultra-high spatial resolution (100x100x200µm) BOLD fMRI enabled the brain-wide mapping of activated visual and vibrissa systems during sensory stimulation in awake mice, including association cortices, e.g. anterior cingulate cortex and retrosplenial cortex, for high order sensory processing. Also, the activation of barrel cortex at 2 s prior to the air-puff indicated a learned anticipation of awake mice under continuous training of the repetitive vibrissa stimulation.

## Introduction

Functional Magnetic Resonance Imaging (fMRI) indirectly measures brain activity via MRI contrast associated with endogenous blood oxygen level dependent (BOLD) signals(1–3). The BOLD contrast was first described by Pauling and Coryell in 1936(4), but it had not been utilized in anesthetized rodent MRI until 1990(1, 2). The power of BOLD-fMRI was later revealed in human brain functional mapping(5–7) and has revolutionized cognitive neuroscience. In contrast to human studies, preclinical fMRI has played a crucial role in method development and validation(3, 8–14). fMRI of anesthetized rodents reduces confounding artifacts due to motion and detects robust BOLD or Cerebral Blood Volume (CBV) signals under various anesthetics(15–29). Recently, the bridging power of preclinical fMRI for basic mechanistic and translational studies has been further exploited given the combination of rodent fMRI with genetic modification tools (e.g., optogenetics, chemogenetics, and genetically encoded biosensors)(8, 30–43). Among the many efforts in anesthetized rodent fMRI, mouse fMRI set a foundation for mechanistic multi- modal imaging given its global mapping scheme in genetic modification models(16, 19, 44, 45), as well as the ability to perform viral transfections to circuit- or cellular-specific targets in transgenic models. However, anesthetics alter brain function during fMRI, preventing accurate interpretation of brain functional changes in awake states(15, 20, 23, 25, 26, 46–52).

Awake mouse fMRI presents itself to provide the most relevant brain functional mapping information for translational cross-scale brain dynamic studies. To immobilize the mouse head during scanning, surgical implantation of headposts has been developed for head-fixation similar to optical imaging schemes(10, 53, 54). In contrast to the fMRI mapping of anesthetized animals, motion-induced artifacts and potential stress-related issues caused by loud gradient noises and micro-vibrations during scanning are major difficulties faced by existing awake mouse fMRI studies(46, 47, 55, 57, 60–63). Previous work has demonstrated that well-planned training procedures could acclimate awake mice during scanning(57, 58, 60–62, 64–66), however; different training paradigms are expected to produce large variability in the functional mapping results(67). One ongoing challenge of awake mouse fMRI is to provide reproducible and high-quality brain functional images with sufficient spatiotemporal resolution and signal-to-noise ratio (SNR) to distinguish functional nuclei of only a few hundred microns in mouse brains. Since increasing spatiotemporal resolution leads to a reduction in SNR of the images, accessing the highest field MRI available, as well as maximizing the efficiency of the Radio Frequency (RF) transceiver signal is critical. Although cryoprobes have been well implemented to boost SNR(10, 47, 68), construction limitations of the superconducting environment constrain the usable space and flexibility to accommodate other imaging/recording modalities(10, 13, 69–74). Implantable coils have been used in animal imaging for over three decades(12, 75–81). Their use gained popularity due to higher SNR and reduction of susceptibility artifacts. The main limitation of implantable coils is the need to surgically implant these coils, adding a degree of invasiveness that MRI usually avoids. However, for typical awake mouse neuroimaging studies, surgical procedures to provide a head-fixation apparatus are routinely practiced. Replacing the conventional head-post for immobilization of the head with an implantable RF coil is critical for achieving high-resolution awake mouse fMRI using ultra-high field MRI, e.g., 14T shown here.

In this present study, we established an awake mouse fMRI platform by applying an implantable RF surface coil, permanently affixed to the head, which simultaneously functioned as a head post for fixation during scanning, minimizing animal motion. This setup allowed us to acquire images with an in-plane spatial resolution of 100µm and 200µm slice thickness. This unique implantable RF coil/headpost scheme simplified the awake mouse training and conditioning for imaging. While there is not currently sufficient evidence to ascertain whether head-fixed training leads to stress-free animals, we observed that a 5-week training scheme resulted in increased eye movements, presenting decreased struggling and freezing behavior indicative of awake mice during scanning. This implanted RF coil scheme also improved B0 homogeneity, as well as effectively eliminated any motion-related loading changes causing B1 variability. Here we successfully mapped activated visual and vibrissa pathways and detected robust BOLD responses in higher-order association cortices, e.g., Anterior Cingulate Area (ACA) with visual stimulation and Ventral Retrosplenial Area (VRA) with vibrissa stimulation in awake mice based on connectivity map projections from the Allen Brain Atlas derived from a Cre-dependent AAV tracing of axonal projections (82). Interestingly, the repetitive vibrissa stimulation paradigm in awake mice has enabled us to detect potential anticipatory learning predicting the onset of stimulation. Our work is a fundamental step towards combining high-resolution fMRI with other modalities to simultaneously record neuronal and microvascular signals throughout brain-wide circuity in awake mice.

## Methods

### Animals

Thirty-eight C57BL/6 mice were used in the current study (weighing between 20 and 30 g) and allocated as follows for each experiment: SNR measurements at 9.4T – sixteen male mice, SNR, visual, & whisker stimulation measurements at 14T – thirteen mice (6F/7M), and random stimulation measurements at 14T – nine mice (4F/5M). Mice were group housed (3-4/cage) under a 12-h light/dark cycle with food and water ad libitum. All animal procedures were conducted in accordance with protocols approved by the Massachusetts General Hospital (MGH) Institutional Animal Care and Use Committee (IACUC), and animals were cared for according to the requirements of the National Research Council’s Guide for the Care and Use of Laboratory Animals.

### Awake mouse fMRI setup

The awake mouse cradle was designed in Blender and 3D printed using a Formlabs 3L 3D printer (Formlabs Inc., Somerville, MA). The design incorporated a sliding track which accepted the PCB chip transceiver circuit to slide in while the mouse was inserted into the cradle. Two transceiver circuit designs were built, a single loop and a figure 8 design. Each one keeps the B1 direction orthogonal to the B0. The single loop allows for full brain coverage at sufficient depths for subcortical investigation. The figure 8 design, due to its smaller coil loops and B1 direction, is limited to precise measurements of shallow brain regions but provides a significant increase in SNR which is beneficial to cortical specific studies which do not have a need to look deeper into subcortical regions but would benefit from a much higher SNR. The single loop or Figure 8 shape RF coils were built to optimize tuning/matching performance when affixed onto the mouse skull. The coils serve to optimize the B0 homogeneity by minimizing the air-tissue interface. Each coil was built to weigh ∼2.5g to minimize the recovery/neck strengthening time of each mouse. The standardized RF coil was acquired from MRIBOT LLC (Malden, MA) using the circuit diagram shown in **supplementary figure 1**.

### Animal Surgery

Mice underwent surgery to affix the RF coil to the head. Animals were anesthetized for surgery using isoflurane. Induction was accomplished using 5% isoflurane and 1L/min of medical air and 0.2L/min additional O2 flow. Animals were maintained at 1.5%-2% isoflurane using respiration rate as a monitor for anesthesia depth. To attach the head coil, mice were affixed in a stereotaxic stage to stabilize the head with ear and bite bars. The scalp was shaved sterilized with ethanol and iodine and an incision was made to expose an area of the scull the size of the RF coil ring. The skull was cleaned of residual tissue and cleaned with 0.3% H2O2 and PBS before being dried. The coil was then positioned over the skull covering the underlying brain. The coil ring was lifted ∼0.3-0.5mm above the surface of the skull to avoid over-loading effects and held in place while a thin layer of cyanoacrylate glue was applied to connect the skull with the coil. Once dried (∼5-8 mins), 2-part dental cement (Stoelting Co., Wood Dale, IL) was mixed and applied to cover the coil and exposed bone paying special note to the base of the coil to firmly secure it and avoid air bubbles and drips toward the eyes. The edges of the skin were then glued to close the surgical site. After the dental cement had fully hardened (∼10 mins), the mouse was released from the stereotaxic stage and received subcutaneous injections of Dexamethasone and Cefazolin. Mice were then allowed to recover in their home cage for at least 1 week to ensure ample neck strengthening had occurred and the mice could walk with normal head posture.

### Animal Training

To acclimate the animals to the MRI environment, each mouse underwent 5-weeks of intermittent habituation procedures to train animals before fMRI experiments by using the following method:

### Training days pre-surgery (Phase 1)

1. in hand mouse handling → 5 min
2. in hand mouse handling → 10 mins

Holder training days post-surgery (after recovery) (Phase 2)

1. secured in holder ◊ 15 mins

a. 10 min pupil recording
2. secured in holder ◊ 30 mins

a. 10 min pupil recording <u>Mock-MRI training days</u> (Phase 3)
3. secured in holder with MRI audio ◊ 30 mins

a. 10 min pupil recording
4. secured in holder with MRI audio ◊ 30 mins

a. 10 min pupil recording
5. secured in holder with MRI audio ◊ 30 mins

a. 10 min pupil recording
6. secured in holder with MRI audio ◊ 30 mins

a. 10 min pupil recording
7. secured in holder with MRI audio ◊ 60 mins

a. 10 min pupil recording
8. secured in holder with MRI audio ◊ 60 mins

a. 10 min pupil recording
9. secured in holder with MRI audio ◊ 60 mins

a. 10 min pupil recording after
10. secured in holder with MRI audio ◊ 60 mins

a. 10 min pupil recording

Training days inside MRI (resting-state & Stimulation) (Phase 4)

1. Real scans (EPI with pupil recording)
2. Real scans (EPI with pupil recording)
3. Real scans (EPI with air puff & pupil recording)
4. Real scans (EPI with air puff & pupil recording)

Following this, data acquisition began, and pupil changes were monitored during scans. During all pupil recordings, animals were secured in the holder and ensured the environment for recording did not allow external light to reach the pupil. Illumination was achieved from a 660nm LED light source (Thorlabs, Inc., Newton, NJ.) delivered via fiber optic cable. Videos were captured at 30fps using a 1/3” CMOS camera and 12mm focal length lens (Tru Components, Chicago, IL)

#### Pupil/Eye fluctuations during training regime

The pupils of awake mice were recorded during training sessions, allowing investigation into eye movements and pupil diameter changes as potential surrogate of stress-related readouts of the animals. The pupil recordings were measured over 12 training days across three-four weeks in Phase 2 to 4 (**Table 1**). In Phase 1, animals were gently held in hands, not head fixed in the cradle. In the second half of phase 4, head-fixed animals were exposed to air puff stimulation inside the MR scanner during echo-planar-imaging (EPI) sequences acquisition. At this stage, animals tended to close their eyes in response to air puff, so pupil measurements were not feasible to be included for data analysis. The increased eye movements were well detected in Phase 3 when animals were exposed to the real MRI acoustic noise in the head-fixed position (located in the RF shielded box attached to the 14T magnet, i.e. the CCM box) (**Supp Fig 2A**). Interestingly, in Phase 4, animals showed significantly reduced eye movements during the first day when animals were positioned inside the MR scanner with EPI scanning, while an increase in eye movement went back to the level of phase 3 in the following training days. The pupil diameter changes were also measured as the function of training days. The power spectral analysis showed ultra-slow pupil dynamic changes with peaked bandwidths less than 0.02Hz. Interestingly, the power of the ultra-slow pupil dynamics also increased as the function of time similar to eye movements, in particular, during phase 3. Meanwhile, power reduction in the first day of in-bore training followed with recovered pupil dynamics in the following days was also observed during phase 4. Although the actual stress of the animals during scanning remains to be further investigated following the 5- week training procedure, the motion-induced image distortion has been dramatically reduced in well-trained animals compared to the start of in-bore training.

**Figure 1.**
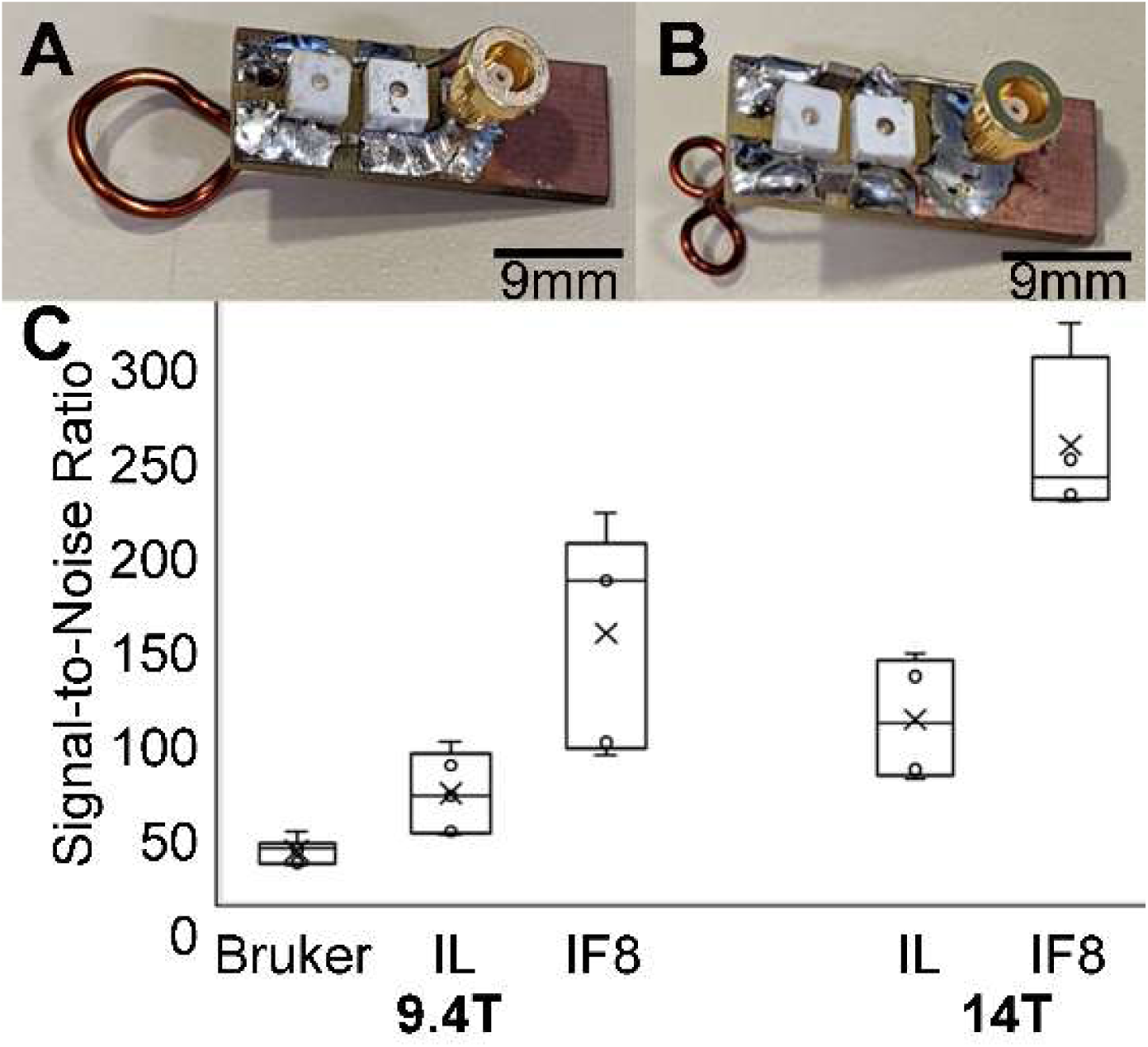
**A** & **B** show representative (unattached) prototype coils in the single loop and figure 8 styles, respectively. **C** presents the cortical specific SNR values calculated by dividing the mean signal of the upper cortex by the standard deviation of the noise to compare between commercial Bruker phased array surface coil, single loop implant and “figure 8” style implants. Bruker ◊ commercial phased array coil, IL ◊ implanted single loop coil, IF8 ◊ implanted “figure 8” coil. The bar graph shows the SNR of anatomical images acquired with different RF coils using the 9.4T scanner 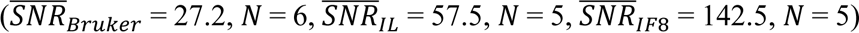 and the 14T scanner 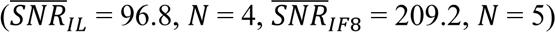

**Figure 2.**
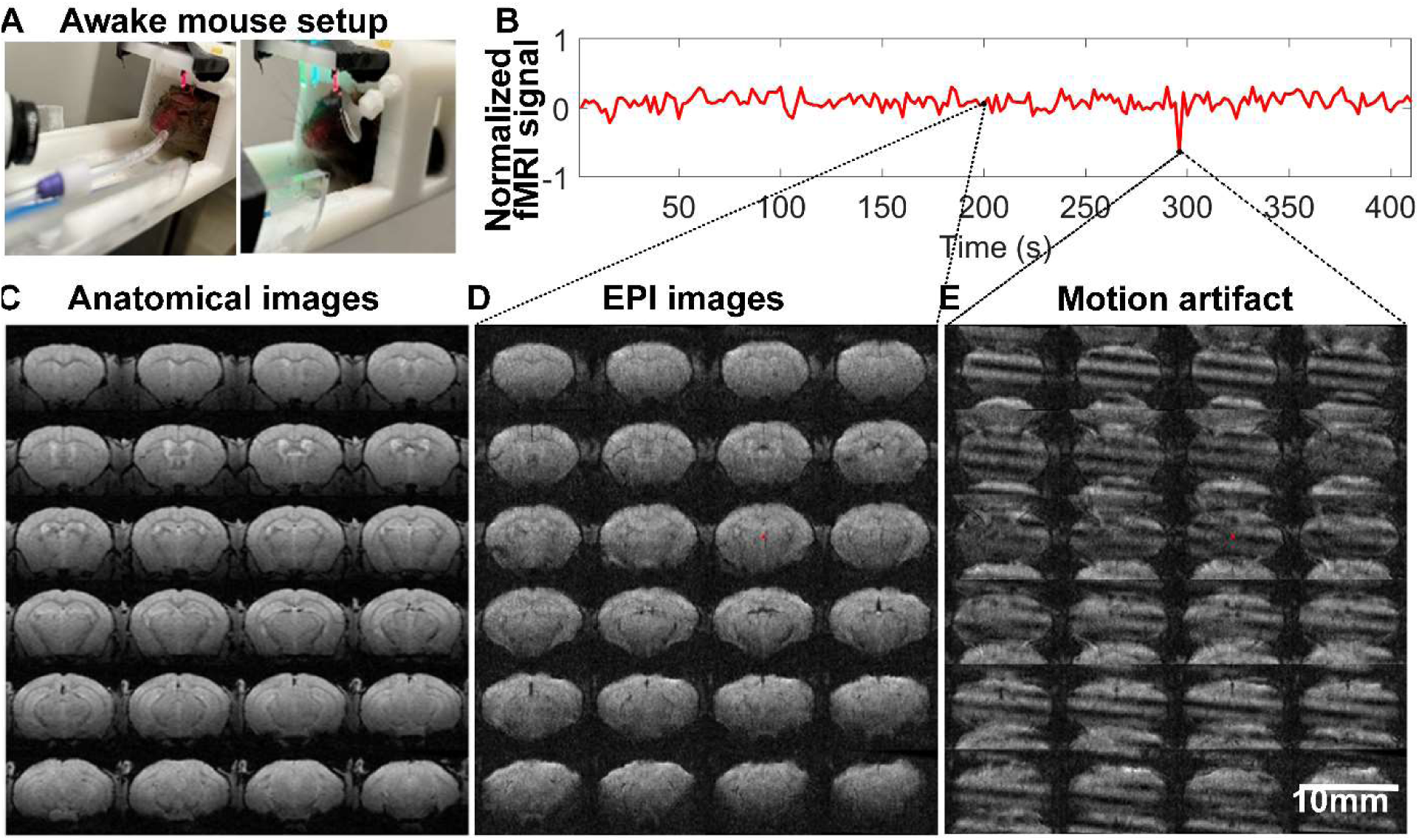
High resolution awake mouse fMRI at 14T. **A**. The awake mouse setup with head-fixed position in a custom-built cradle for visual and vibrissa stimulation. **B.** The representative fMRI time course of an awake mouse based on raw image data acquired from high-resolution EPI, enabling the trace of motion-induced artifacts. **C**. The anatomical MRI images (FLASH) acquired from one representative awake mouse, showing minimal susceptibility and whole brain coverage from the implanted surface coil. **D**. The raw EPI fMRI image with same spatial resolution as the anatomical FLASH image. **E**. The snapshot of the distorted images due to motion of the awake mouse during scanning (Supp Movie 2 shows the video of motion-induced artifacts throughout the fMRI trial).

**Table 1:** Table showing the training paradigm for each of the 4 phases of training. rs ◊ resting state fMRI, stim ◊ whisker stimulation fMRI.

| Training Day of Phase | Phase 1 (hold in hand) | Phase 2 (holder + pupil) | Phase 3 (Mock-MRI + pupil) | Phase 4 (EPI + pupil) |
| --- | --- | --- | --- | --- |
| 1 | 5 mins | 15 mins | 30 mins | 60 mins (rs) |
| 2 | 10 mins | 30 mins | 30 mins | 60 mins (rs) |
| 3 |  |  | 30 mins | 60 mins (stim) |
| 4 |  |  | 30 mins | 60 mins (stim) |
| 5 |  |  | 60 mins |  |
| 6 |  |  | 60 mins |  |
| 7 |  |  | 60 mins |  |
| 8 |  |  | 60 mins |  |

### Anesthesia Regiment for MRI measurements of SNR

While acquiring images to measure SNR improvements, all animals were anesthetized for the duration of MR scanning. Mice were induced using 5% isoflurane in medical air and maintained with 1.0-2.0% isoflurane, adjusted to retain stable physiological conditions while in the magnet. The gas mixture was supplied through the hollow bite bar directly to the mouth and nose of the animal at a rate of 1.0L/min. Animals were anesthetized to minimize artifacts associated with motion. Physiological monitoring of the animal was performed through the integration of a Small Animal Monitoring and Gating System (Model 1030, SA Instruments, Inc., Stony Brook, NY) capable of recording respiration, body temperature, electrocardiogram, and other parameters. The animal’s breathing rate was continuously monitored and recorded during scanning using a pressure-sensitive sensor-pad and maintained between 50-80 breaths/min. Animals were kept at a constant temperature of 37°C in the MRI scanner by means of blowing warm air through the bore and recorded using a rectal thermometer probe.

### MRI methods

^1^H MRI data was acquired using the 14T and 9.4T horizontal MRI scanners (Magnex Sci, UK) located at the Athinoula A. Martinos Center for Biomedical Imaging in Boston, MA. The 14T magnet is equipped with a Bruker Avance Neo Console (Bruker-Biospin, Billerica, MA) and is operated using ParaVision 360 V.3.3. A microimaging gradient system (Resonance Research, Inc., Billerica, MA) provides a peak gradient strength of 1.2T/m over a 60-mm diameter. The 9.4T scanner is equipped with a Bruker Avance III HD Console (Bruker-Biospin, Billerica, MA) and is operated using ParaVision 6. A dual microimaging gradient system comprises a Bruker gradient coil capable of 44 G/cm, and a Resonance Research gradient insert capable of 150 G/cm.

#### SNR Measurements

^1^H MRI data for SNR measurements were acquired on 9.4T (400MHz) and 14T (600MHZ) scanners using the following parameters for both systems: TE/TR = 3ms/475ms, flip angle = 30°, and 4 averages for an approximate acquisition time of 4.5 minutes.

9.4T scanner was only used to show SNR improvements from the implantable coils. The BOLD fMRI data were collected only at 14T due to the much-improved SNR available and were collected solely in awake mice to investigate signal associated with the awake functional connectivity. Furthermore, the figure 8 coils were only used to show the SNR improvement due to the coil design for cortical measurements.

#### fMRI BOLD Imaging

Multislice 2D gradient-echo Echo Planar Imaging (EPI) was used to acquire fMRI BOLD data from the awake animals with the following parameters: TE/TR = 7ms/1s, segments = 2, bandwidth = 277,777Hz, 100µm x 100µm in plane resolution with a 200µm slice thickness, 36 slices, 205 repetitions for an acquisition time of 6 minutes 50 seconds.

#### Anatomical Imaging

^1^H MRI data for anatomical registration data were acquired using a multi-slice T1-weighted 2D gradient echo Fast Low Angle SHot (FLASH) sequence with the same parameters of the SNR measurement scans except the resolution was adjusted to match the BOLD data at 100µm x 100µm x 200µm resolution.

### Stimulation method/paradigm

The visual and vibrissa stimulation block paradigm was designed as follows: 5 baseline scans, 1 stimulation trigger scan, 19 inter-stimulation scans, and 10 epochs. The visual stimulation used 2 different wavelengths of light: 530nm and 490nm, which flashed at 5Hz and 5.1Hz, respectively, for 8 seconds with a 20ms “on” time of each illumination. The whisker air puff stimulation used the same block design as the visual stimulation but stimulated with a 10ms puff duration and an 8Hz firing rate for 8 seconds. Due to the use of 2 segments for these experiments, the effective TR was 2s. Therefore, the stimulation duration for both visual and vibrissa experiments resulted in 4 consecutive scans being included in the “on” stimulation period and 16 consecutive scans being included in the “rest” period (**Supp Fig 3**). The random vibrissa stimulation paradigm used the same 10ms air puff at 8Hz for 8 seconds but randomized the “rest” duration. “Rest” durations were 12s, 22s, 32s, and 42s and randomized in 3 different sequences maintaining a scan duration of 205 TRs for each experiment.

**Figure 3.**
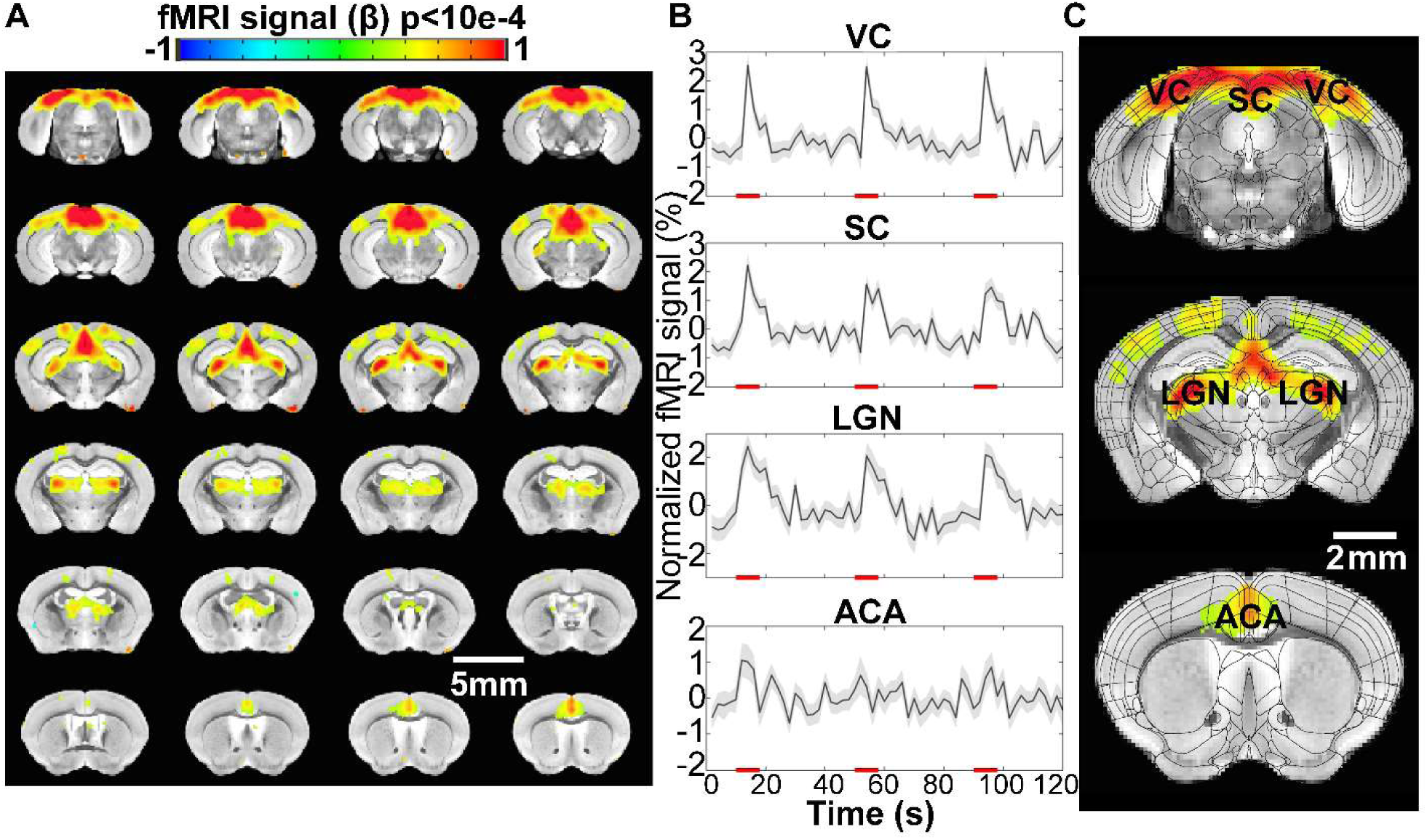
Visual stimulation-evoked high-resolution fMRI of awake mice. **A.** The brain-wide functional maps of awake mice show strong positive BOLD activation in the visual cortex (VC), lateral geniculate nucleus (LGN), superior colliculus (SC), and anterior cingulate area (ACA)) based on the group analysis. **B.** The averaged time course of the ROIs derived from the Allan brain atlas, demonstrating an evoked positive BOLD signal changes upon the 8s visual stimulation (5Hz 530nm and 5.1Hz 490nm 20ms light pluses). Each graph displays the average of 162 sets of 3 stimulation epochs. Shaded regions represent standard error. Red lines represent the 8s stimulation duration. **C.** Functional maps overlain with the brain atlas to highlight the activated brains regions: VC, SC, LGN, and ACA. (N = 13 (6F/7M)).

### Processing/Analysis methods (AFNI and MATLAB))

SNR was computed by dividing mean signal over the standard deviation of the noise. SNR line profile signal data was collected using Amira software (Thermo Fischer Scientific Inc., Waltham, MA). fMRI data was processed using Analysis of Functional Neuroimages (AFNI)(83, 84). Bruker 2dseq images of the EPI and FLASH scans were converted to AFNI format using ‘to3d’ before masking and aligning the dataset to a template.

To process the high resolution stimulated BOLD response from the visual and vibrissa stimulation paradigms, we developed a processing pipeline (**Supp Fig 4**). For each experiment, FLASH data were averaged for anatomical localization and the EPI scans were time averaged before registration. The time-averaged EPI was then registered to the FLASH and then to the Australian Mouse Brain Mapping Consortium (AMBMC) atlas(85) where a mask was generated. Each time series for each experiment was concatenated so each experiment contains one long time series dataset using the ‘3dTcat’ command. Data were then despiked before each EPI time point was registered, via a 6-degree transformation, to the atlas using the ‘volreg’ command after which the previously generated mask was applied. The ‘blur’ command was used to smooth the newly transformed data before it was scaled and underwent a linear regression. All concatenated data was then split and summed, per each experimental study, to undergo motion correction and outlier removal. The corrected data was then summed and averaged with the remaining processed data to generate a single time series across all experiments. A clustering threshold was set at 100 voxels and the Pearson correlation values were limited to p≤0.01 (corrected) with estimated false discover rate at q=0.00078. For the random stimulation design, the 3 runs were concatenated into a single time series each ensuring they all followed the same series of random timings.

**Figure 4.**
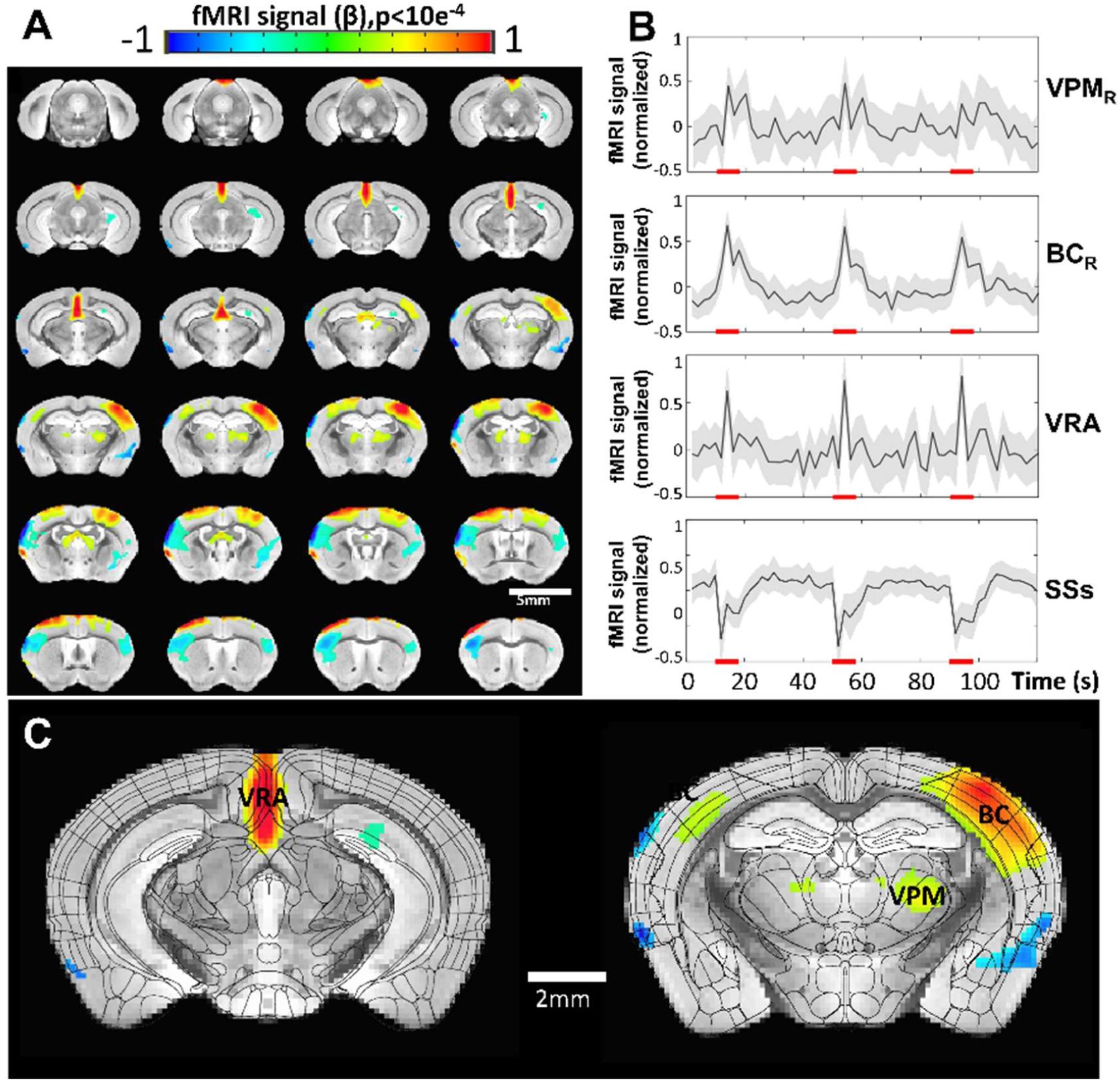
Vibrissa stimulation-evoked high-resolution fMRI of awake mice. **A**. The brain-wide functional maps of awake mice show the strong positive BOLD activation in the contralateral barrel cortex (BC) and ventral posteromedial nucleus (VPM) and posterior thalamic nucleus (PO). Positive BOLD signals are also detected at the motor cortex (MC) and the ventral retrosplenial area (VRA), as well as at the ipsilateral BC and thalamic nuclei. Negative BOLD signals are detected in supplementary somatosensory areas (SSs) (including nose and mouth) as well as part of the caudoputamen. **B.** The averaged time course based on the brain atlas ROIs for VMP, BC, and VRA, demonstrating positive BOLD signal changes upon the 8s air-puff vibrissa stimulation (8Hz, 10ms). Averaged time course of the SSs ROI shows negative BOLD signal changes. Each graph displays the average of 279 sets of 3 stimulation epochs. Shaded regions represent standard error. Red lines represent the 8s stimulation duration. **C.** The functional maps are overlain with the brain atlas to highlight the activated vibrissa thalamocortical pathway (VPM→BC) and the VRA in awake mice. (N = 13 (6F/7M)).

## Results

### Development and efficiency validation of implantable RF coils to boost SNR

We have developed an implantable RF coil which effectively boosted the SNR in ultra-high field MRI. Here we compared two prototypes: a simple single loop coil design and a “figure 8” coil design. These coils were used to check SNR in anatomical data of anesthetized mice at 9.4T and 14T. **Figure 1** shows examples of the prototyped RF coils (**Figure 1A**). The acquired SNR values for each prototype design are shown in **Figure 1B**. Here, we used a commercial 4 phase-array coil (400MHz for 9.4T) as a control to compare with the implantable RF coils. The single loop implantable coils improved SNR over 100% compared the commercial option while the figure 8 style showed a more than 5 times increase at 9.4T in the cortical regions. The SNR along the dorsal-ventral axis was plotted to compare the B1-field sensitivity of the single-loop and figure 8 RF coils in comparison with the phase-array coil at 9.4T, showing significantly increased SNR up to 4 mm depth (**Figure 1B**). Moving up to 14T, the SNR improvements increased proportionally as a factor of field strength(86). This improved SNR allows for high spatial resolution fMRI studies of awake mice. The single loop coil tuned to 14T (600MHz) was used for functional data collection in the manuscript. The figure 8 coil was only used as part of development to show the improvement of the coil design for cortical MR signal measurements.

### Awake mouse fMRI with visual stimulation

The RF coil was implanted on the mouse skull to serve as an attachment for head fixation during awake mouse fMRI at 14T (**Figure 2A**). The awake mouse fMRI setup was designed using a 3D printed cradle incorporated a sliding track which enabled the printed circuit board (PCB) chip mounted on the mouse head to slide through. The PCB chip was then fixed in place at the end of the cradle using friction screws (**Supp Movie 1**). Once the mouse was fixed in the animal cradle, either a mirror or air tube was positioned for pupillometry recording or vibrissa stimulation, respectively. Additionally, an MRI-compatible camera was incorporated to record the pupil dynamic changes and whisking behavior of awake mice during scanning. One key feature of the awake mouse fMRI setup is the plug and play capability for scanning.

This awake mouse fMRI setup enabled high-resolution echo planar imaging (EPI) data acquisition at 100x100x200um spatial resolution with a 2s effective repetition time (TR). The EPI-based T2* images acquired from head-fixed awake mice show little air-tissue interface-induced image distortion with the same spatial resolution as anatomical images (**Figure 2C, D**). Motion artifacts were detected at some time points of the fMRI time course, presenting large EPI image distortions (**Figure 3E, Supp Movie 2**), but can be removed using a censoring function during data analysis (**Supp Fig 4**). These results have shown that the multi-slice 2D EPI enables brain-wide functional mapping of awake mice.

To map the brain function of awake mice with this high-resolution fMRI method, we first introduced a visual stimulation paradigm. Based on a block-design regression analysis, we detected robust BOLD responses. Brain-wide functional maps using the visual stimulation paradigm were seen with activated areas highlighted along the visual pathways (**Figure 3**). These areas included the visual cortex (VC), superior colliculus (SC), lateral geniculate nucleus (LGN), and association cortex in the anterior cingulate area (ACA). The ROI-specific localization was well characterized by overlapping the brain atlas and functional maps (**Figure 3C**). The ROI-based time courses demonstrated robust BOLD responses detected in awake mice (**Figure 3B**).

### Awake mouse fMRI with vibrissa stimulation

In contrast to a visual sensation, awake mice may flinch due to the sudden physical vibrissa stimulation causing severe motion artifacts during scanning. Prolonged training was needed to reduce motion artifacts during air-puff stimulation as shown in **Supp Movie 2** allowing for high-resolution fMRI of awake mice. The activated barrel cortex (BC) and ventroposterior medial nucleus (VPM) were seen related to stimulation of the contralateral whisker pad (time courses in **Figure 4B**, **Supp Fig 5**). Brain-wide functional maps also showed activation in the motor cortex and a small portion of the ipsilateral BC. These results demonstrated the importance of distinguishing BOLD activation between external stimulation and voluntary movements while also confirming the feasibility to map brain-wide brain activations in awake behaving mice with 14T fMRI.

**Figure 5.**
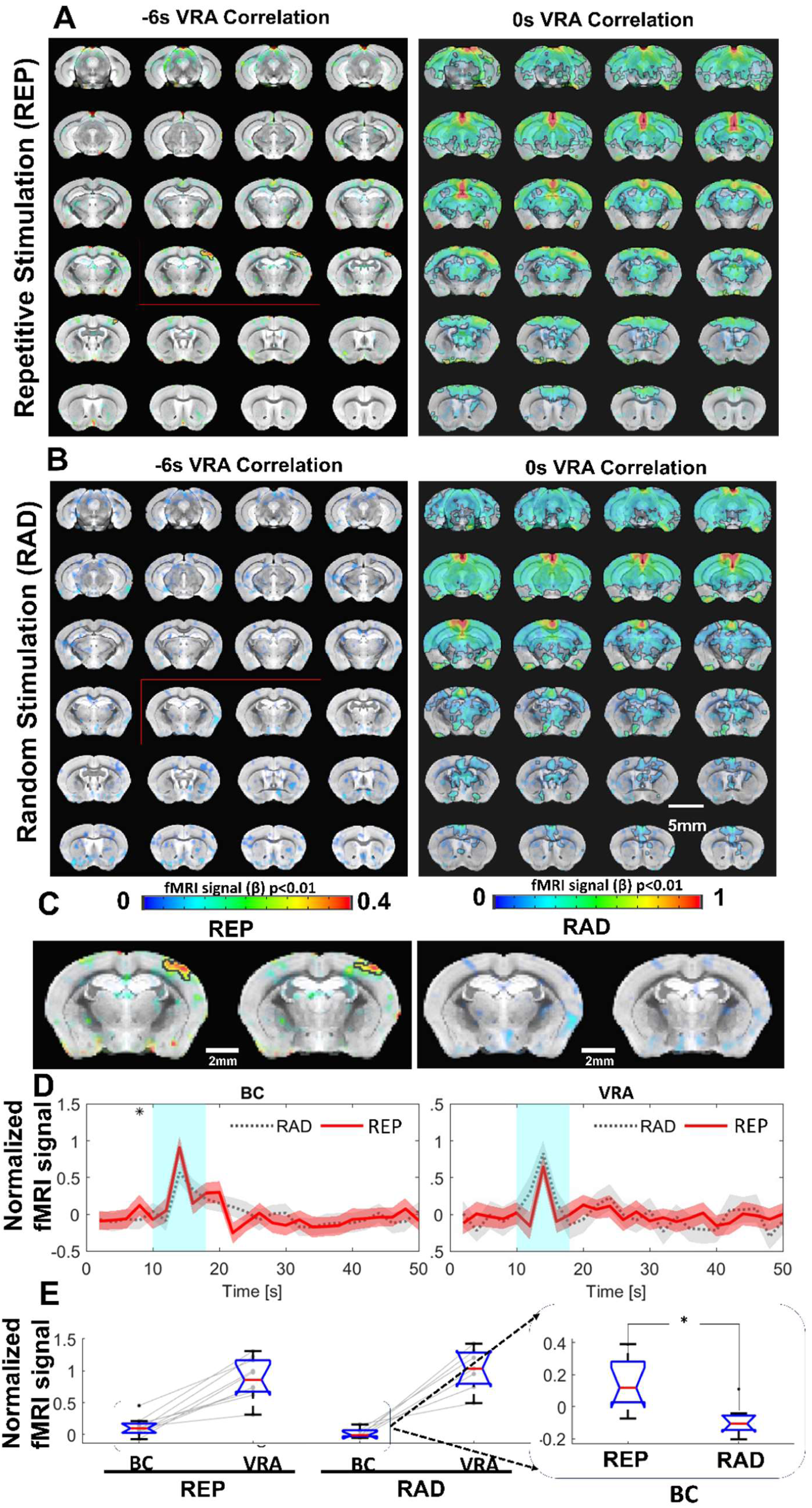
VRA-based brain-wide correlation maps at different time shifts. **A**. The VRA-based correlation maps at -6s and 0s time shifts of awake mice with repetitive stimulation (REP). The strong correlation in the contralateral BC is shown in the correlation map at the -6s time shift (red box). **B**. The VRA-based correlation maps at -6s and 0s time shifts of awake mice with randomized stimulation (RAD). No correlation is detected in the contralateral BC at the -6s time shift (red box). **C.** The enlarged images from the -6s time shift correlation maps of REP and RAD groups, demonstrating the strong correlation patterns located at the contralateral BC only in the REP group. **D**. The averaged time course from both contralateral BC and VRA of REP and RAD groups, showing that early positive BOLD signals detected at 2 s prior to the stimulation in contralateral BC of the REP group and no significance difference detected in VRA. **E**. The bar graph presents the mean BOLD signals of contralateral BC at 2s prior to stimulation time point and peak signals of VRA in REP and RAD groups. The inset is the expanded bar graph to show the significantly higher BOLD signals detected in the contralateral BC at 2 s prior to stimulation in REP group (p=0.015, REP graph displays the average of 930 stimulation epochs, RAD graph displays the average of 240 stimulation epochs). (N = 9 (4F/5M)).

### Prediction-related barrel cortical activity to patterned air-puff in awake mice

An interesting observation from the vibrissa stimulation was the activated VRA. The VRA only showed brief responses to air puff in contrast to the typical duration of hemodynamic responses observed in the BC and VPM. This presents a good landmark for studying higher level processing of vibrissa sensation. Voxel-wise cross correlation analysis was performed based on the VRA-specific fMRI dynamic changes. At a zero time-shift (map developed from peak BOLD response), the VRA is strongly correlated with the hippocampus, cingulate cortex, and central thalamic regions (**Figure 5A**). Nevertheless, at a -6s time-shift (i.e., 2s before stimulation onset), stronger correlation was observed at the contralateral BC, indicating anticipation of the repetitive air-puff in the block design (**Figure 5A**). To validate that this early BC activation was caused by learned anticipation of the time-fixed repetitive air-puff stimulation, we also analyzed the VRA- specific cross correlation in a control group using a randomized stimulation paradigm. Although VRA remained strongly coupled with the other association cortices and subcortical regions at the zero time-shift, no correlation was observed from the contralateral BC at the -6s time-shift (**Figures 5B**). The fMRI time course analysis from the contralateral BC also showed increased BOLD responses before the air-puff in the block design group, but not the randomized control group (**Figure 5C**). Quantitative analysis showed a significantly higher BOLD signal two seconds before the air puff stimulation in the standard block design group when compared with the randomized group (**Figure 5D**). It should be noted that VRA responses between the two groups were similar, further confirming the anticipation-related early BC activation to repetitive air-puff stimulation.

## Discussion

In this study, we designed and implemented implantable RF coils for awake mouse fMRI, which also served as a headposts during data acquisition. Our design, based on previously published cable-free (inductive) RF coils(81), offered an easier pre-scan setup by eliminating the need to localize and secure the pickup coil for inductive coupling optimization. And while this current design showed reduced freedom for animal movement, implanted coils offer more stable sample loading and reduce the B0 offset when compared to the previous version. This was also true when comparing to conventional RF coils as the motion of the animal would alter the loading and cause B1 field variability during fMRI scanning.

### Technical considerations with awake mouse fMRI at 14T

A few important factors should be considered to improve data quality using the implanted RF coils described in the present study. The first is animal motion. As this design was used for awake and minimally restrained mice, the animals would eventually move to adjust themselves (scratching, grooming, teeth grinding, etc.) during the scan. This will affect B0 homogeneity and can cause ghosting if severe enough. This can be minimized through acclimation training which will also reduce unwanted stress. Other studies have animal restraint mechanisms that seek to restrain the body of the animal(46, 47, 58, 87) but has the potential to cause unwanted stress which can affect the desired fMRI signals. Furthermore, B1 variability was present through motion as well due to the current design of the coil. As the RF circuit chip sits above the animal’s neck, body movement could alter the loading of the circuit, inducing B1 artifacts through lifting or dropping the body towards or away from the circuit chip. Again, these artifacts can be minimized through proper training and stress reduction which was well accomplished for our study through the design and training method(60). Here we see that while animal motion still exists (**Supp Fig 6**) large struggling movements are minimized after the training method. Still, mice have a thin skull, which leads to the air-tissue interface being a non-negligible factor at ultra-high fields (e.g. 14T). Therefore, the coil implantation shown here has reduced this source of inhomogeneity (**Supp Fig 2 & 6**) and allowed for a consistent and stable shim. By implanting the coil on the surface of the skull, we can achieve a significantly higher SNR which can be comparable to cryoprobe designs at close distances. Moreso, this is the case with the figure 8 coil shown in this study which gives a 5-times increase in SNR over the standard commercially available 4 array mouse head coil for 9.4T (both operating at room temperature) **(Figure 1**). This improvement allows much higher spatial resolution in awake mouse fMRI, at a two nanoliter voxel volume, compared to contemporary efforts in human brain mapping at sub-millimeter resolution (0.5-0.8mm isotropic), a difference of two orders of magnitude(88–91).

### The consideration of stress issues of awake mouse fMRI

Despite the extensive training procedure of the present work in comparison to the existing awake mouse fMRI studies (training strategies for awake mice fMRI have been reviewed by Mandino et al. to show the overall training duration of existing studies (67)), stress remains a confounding factor for the brain functional mapping in head-fixed mice. During animal training, we have measured both pupil dynamic and eye motion features from training sessions, both of which could be considered as potential surrogate of the stress levels of animals. It should be noted that stress may be related to increased frequency of eye blinking or twitching movements in human subjects(92–94). However, the eyeblink of head-fixed mice has been used for behavioral conditioning to investigate motor learning in normal behaving mice(95–97). Importantly, head-fixed mouse studies have shown that eye movements are significantly reduced compared to the free-moving mice(98). The increased eye movement during acclimation process would indicate an alleviated stress level of the head-fixed mice in our cases. Meanwhile, stress-related pupillary dilation could dominate the pupil dynamics at the early phase of training(63). We have observed a gradually increased pupil dynamic power spectrum at the ultra-slow frequency during phase 3, presenting the alleviated stress-related pupil dilation but recovered pupil dynamics to other factors, including arousal, locomotion, startles, etc. in normal behaving mice. Nevertheless, a recent study(99) shows that the corticosterone concentration in the blood samples of head-fixed mice is significantly reduced on Day 25 following the training but remains higher than in the control mice. Also, the time-dependent changes of stress level during scanning could further confound the functional mapping results if longer than one hour. Thus, the impact of stress on brain functional mapping with awake mouse fMRI would need further investigation, of which the stress-related functional changes should not be neglected from the existing studies.

### Brain-wide functional mapping with visual and vibrissa stimulation

There are fMRI studies investigating the visual system in both anesthetized and awake mice (56, 63, 100–103). In contrast to brain activation patterns at the VC, SC, and LGN(56), robust ACA activation was also detected for awake mouse fMRI in this study (**Fig 3**). Since ACA has been closely involved in pupil dynamics, as well as arousal state regulation(104–106), the mapping of the ACA in awake mice during visual stimulation provides a meaningful way to validate the conscious state of mice during scanning. Similarly, there are extensive rodent fMRI studies of vibrissa stimulation (44, 107–111). In contrast to the anesthetized state, this awake mouse fMRI detected not only activated contralateral BC and VPM, but also spread activation in the motor cortex, and small portion of the ipsilateral BC with positive BOLD signals. Although the air-puff stimulation was set and verified to deflect the whiskers of chosen side, videos of the mouse during scanning show that active bilateral whisking could be initiated upon air-puff. This could lead to bilateral activation of the motor cortex and the ipsilateral BC. Furthermore, studies have been performed to understand the transcallosal activity-mediated excitatory/inhibitory circuits by both fMRI and optical imaging(81, 112–116). The potential transcallosal mediation of the negative BOLD signal detected in the superficial cortical area near BC will need to be further investigated. Also, these negative BOLD signals were detected across a large brain area, which is consistent with astrocyte-mediated negative BOLD during brain state changes reported in anesthetized rats(29) and eye open/close-coupled arousal changes in unanesthetized monkeys(117). Although astrocytic Ca^2+^ transients coincide with positive BOLD responses in the activated cortical areas, which align with the neurovascular coupling (NVC) mechanism(118), there is emerging evidence to show that astrocytic Ca^2+^ transients are coupled with both positive and negative BOLD responses in anesthetized rats(29) and awake mice(119). An intriguing observation is that cortex-wide negative BOLD signals coupled with the spontaneous astrocytic Ca^2+^ transients could co- exist with the positive BOLD signal detected at the activated cortex. Studies have shown that astrocytes are involved in regulating brain state changes(120), in particular, during locomotion(121), and startle responses(122). These brain state-dependent global negative BOLD responses are also related to the arousal changes of both non-human primates(117) and human subjects(123). The established awake mouse fMRI platform with ultra-high spatial resolution will enable the brain-wide activity mapping of the functional nuclei contributing to the brain state changes of head-fixed awake mice in future studies.

Interestingly, vibrissa stimulation also led to robust VRA activation in awake mice(124). VRA serves as one of the major nodes of the default mode network (DMN) across different species(125–129). The vibrissa stimulation-evoked VRA activation suggests the higher-level cortical function contribute to vibrissa sensory processing in awake mice.

### VRA-coupled pre-stimulus BC activation in awake mice as a sign of anticipation

There are extensive studies investigating brain activation responsible for anticipation with fMRI and electrophysiological recordings(130–134). In contrast to the reward anticipation or audiovisual anticipation of naturalistic music and movie clips that demand more complex cognitive processing(135–138), the repetitive air-puff stimulation delivered during head-fixed training for fMRI studies could serve as a simple paradigm to process the anticipatory responses in awake mice. Based on cross-correlation analysis with evoked VRA BOLD responses, the strongest correlation with the BC was detected from 6s lag-time based correlation maps, showing a positive BOLD signal at a time point 2 sec prior to stimulus onset (**Figure 5**). This anticipatory BC response was not detected when the air-puff stimulation paradigm was randomized in another group of mice. VRA is known to be involved in prediction(139–141) and has been coupled with temporal prediction in rodents(140, 142), as well as navigation efficiency involving spatial reference cues(140–142). Additionally, external somatosensory cues (e.g. the air puff or brushing of whiskers) are an important factor when investigating prediction processing(143–147). Previous work has shown that prediction of external stimulation will cause a hemodynamic response even in the absence of a stimulus(134, 148). In our study, we show that after continued regularly spaced stimulation, early somatosensory hemodynamic responses begin to have a significant impact seen in the averaged BOLD response time course. These anticipatory hemodynamic responses are a result of the continuous training for mice experiencing months of repetitive stimulation. The increased BOLD signal in the BC before the stimulus onset shows strong cross-correlation to the VRA activation, but VRA activation is not dependent on the pre-stimulus activation in the BC. This can be seen though the comparable VRA BOLD responses between repetitive and randomized air-puff stimulation paradigms (**Figure 5C**). This result indicates that VRA response mediates external sensory perception and may serve as a key association cortical area for the processing of the anticipated vibrissa signals but is not solely dependent on the prediction of incoming stimulus.

## Data Availability

Data is available for download from OpenNeuro:

Whisker (https://doi.org/10.18112/openneuro.ds005496.v1.0.1), Visual (https://doi.org/10.18112/openneuro.ds005497.v1.0.0) and Zenodo:

SNR Line Profile Data & Data Processing Scripts: (https://zenodo.org/doi/10.5281/zenodo.13821455)

## Acknowledgements

The research benefited from funding from the NIH Brain Initiative grants (RF1NS113278, RF1NS124778, R01NS122904, R01NS120594, and R21NS121642), U19 Cooperative Agreement Grant (U19NS123717), S10 instrument grants (S10OD028616 and S10RR025563) to the Massachusetts General Hospital/Harvard-MIT Program in Health Sciences and Technology Martinos Center, and NSF CBET grant (2123970).

## Author Contributions

DH, XL, XY designed the research. DH, XL, ZX, BZ, AL, XY performed the research. DH, XL, SC, XAZ, AM, YJ analyzed data. ZX, XAZ performed surgeries. DH, BZ, AL, AM built coils. DH, XL, BZ, AL designed animal cradle. DH, XL, AD, XY wrote the paper.

## Supplementary materials and Figures

**Supp Fig 1.**
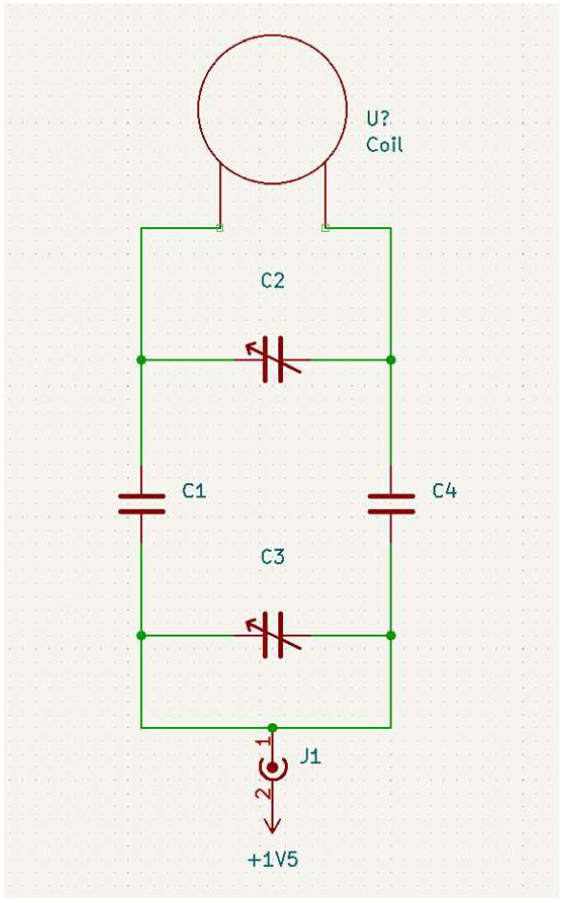
Schematic of circuit diagram of coil designed for ^1^H imaging at 600MHz. C1 & C4 are 2.2pF capacitors. C2 is a 0.6-2.5pF trimmer. C3 is a 5-18pF trimmer.

**Supp Fig 2.**
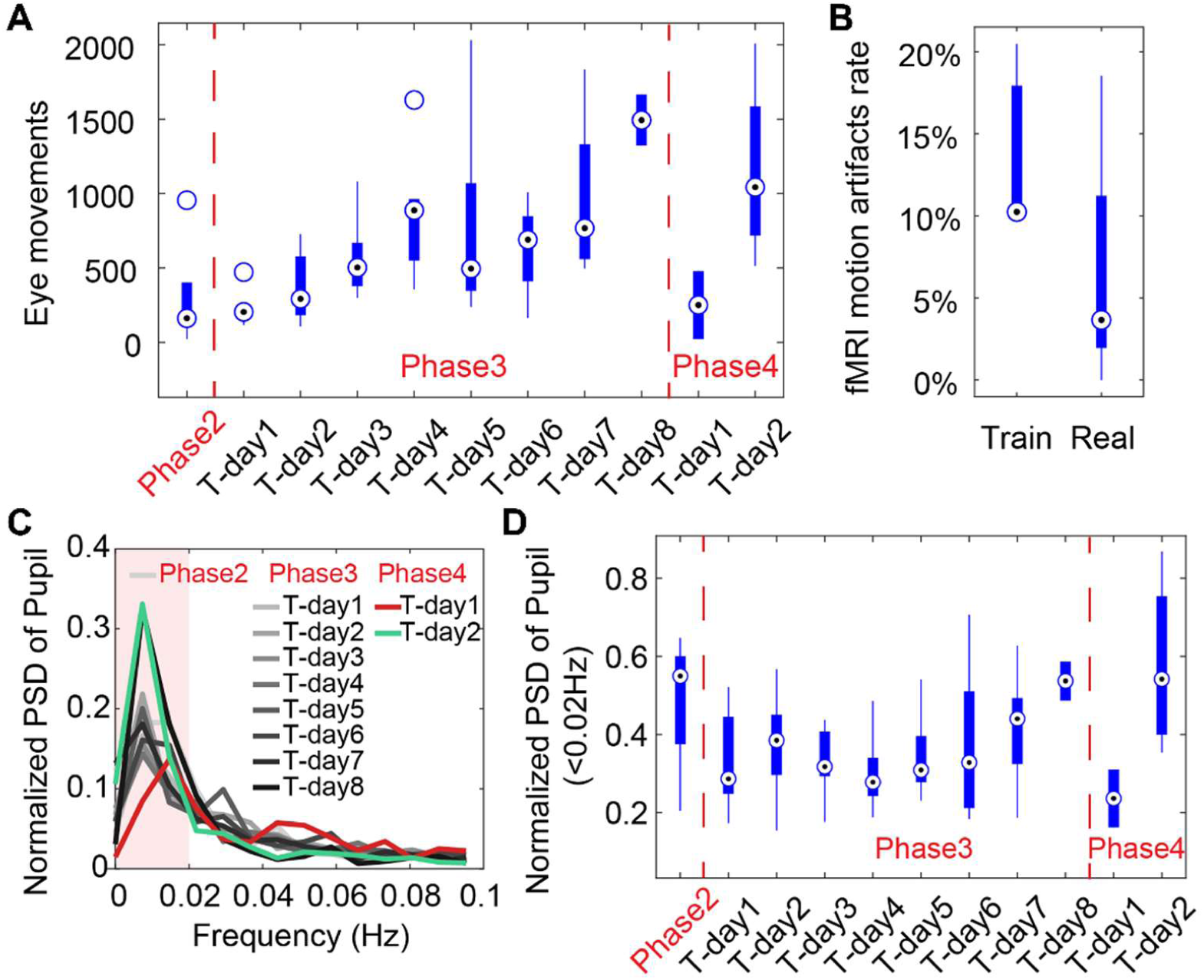
**A** Graph showing eye movements over 10-minute duration of training days showing an increase in eye movements over the training duration indicating the animals are becoming more relaxed in the holder while being exposed to MRI sounds. The drop in eye movements at Phase 4 Training Day 1 is caused by putting the animals into the MRI for the final training. The following day at Phase 4 Training Day 2, the eye movements have increased back to the level before being inserted into the scanner. Phase 2 incorporated the 2 days animals were put in the holder without the mock-MRI environment. Phase 3 includes each day of training in a mock-MRI environment. Phase 4 showcases eye movements acquired inside the MRI during resting state data acquisition. **B.** Graph of the ratio of detectable motion during scanning +1 or 2 days after training (train) and after 1 month of scanning (real) showing a non-significant decrease in the detectable motion indicating the effectiveness of the training. **C.** Graph of the normalized Power Spectrum Density (PSD) showing the increase of low frequency pupil oscillations from eye in each phase. **D.** Averaged normalized PSD graph of pupil oscillation frequencies below 0.02 HZ as maximally indicated in **C** showing late stage increases in mock-MRI and EPI-MRI phases, e.g. phase 3 and phase 4.

**Supp Fig 3.**
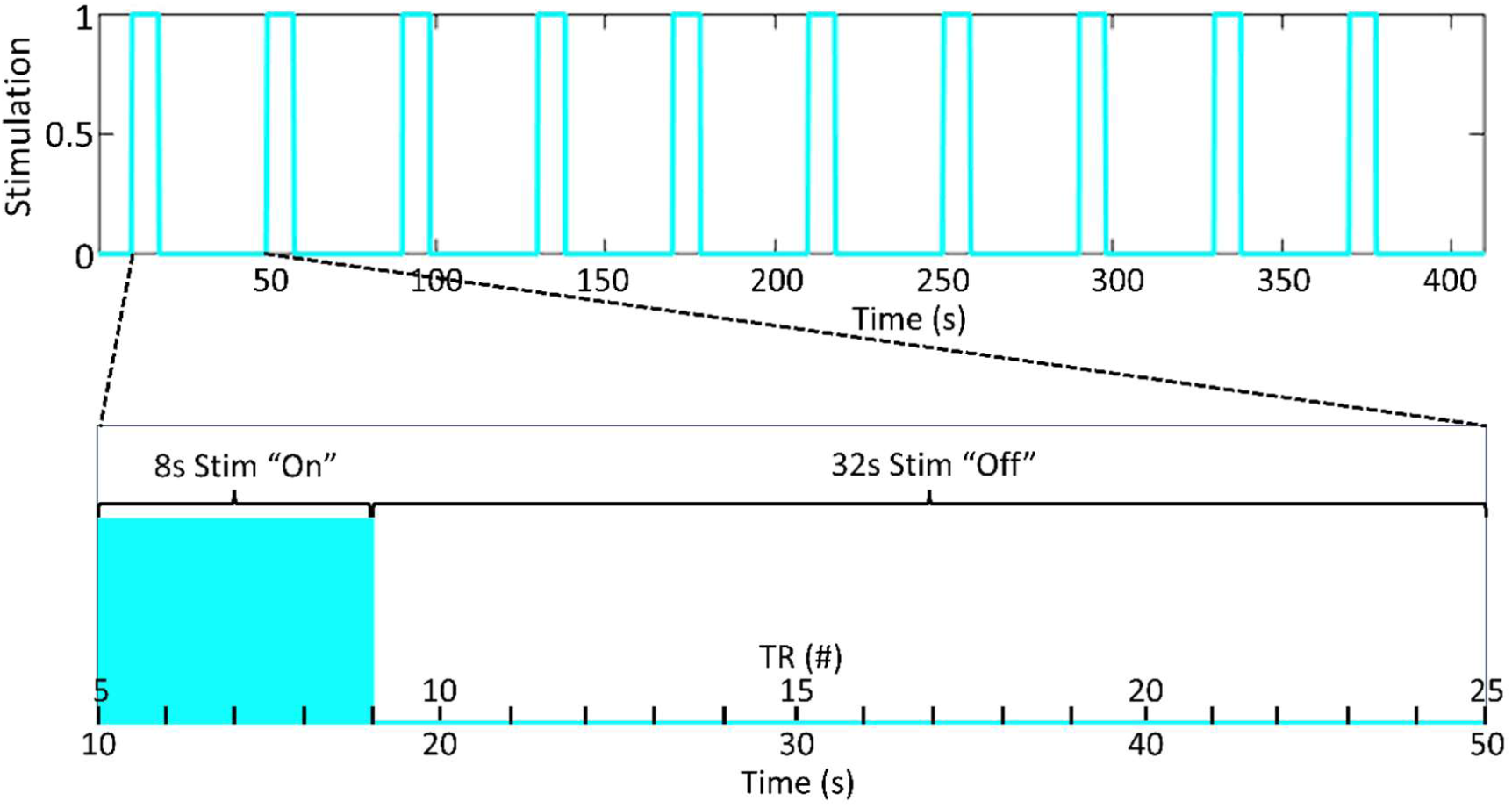
Figure showing stimulation paradigm in blue. Each “TR period” is 2 seconds. The stimulation is repeated 10 times over the 410 second acquisition period which includes a 10 second baseline acquisition period before the first stimulation. This paradigm results in 4 “TR periods” occurring during the stim on phase and 16 “TR periods” occurring during the stim off phase. Any stimulation design can be used for the “on” duration.

**Supp Fig 4.**
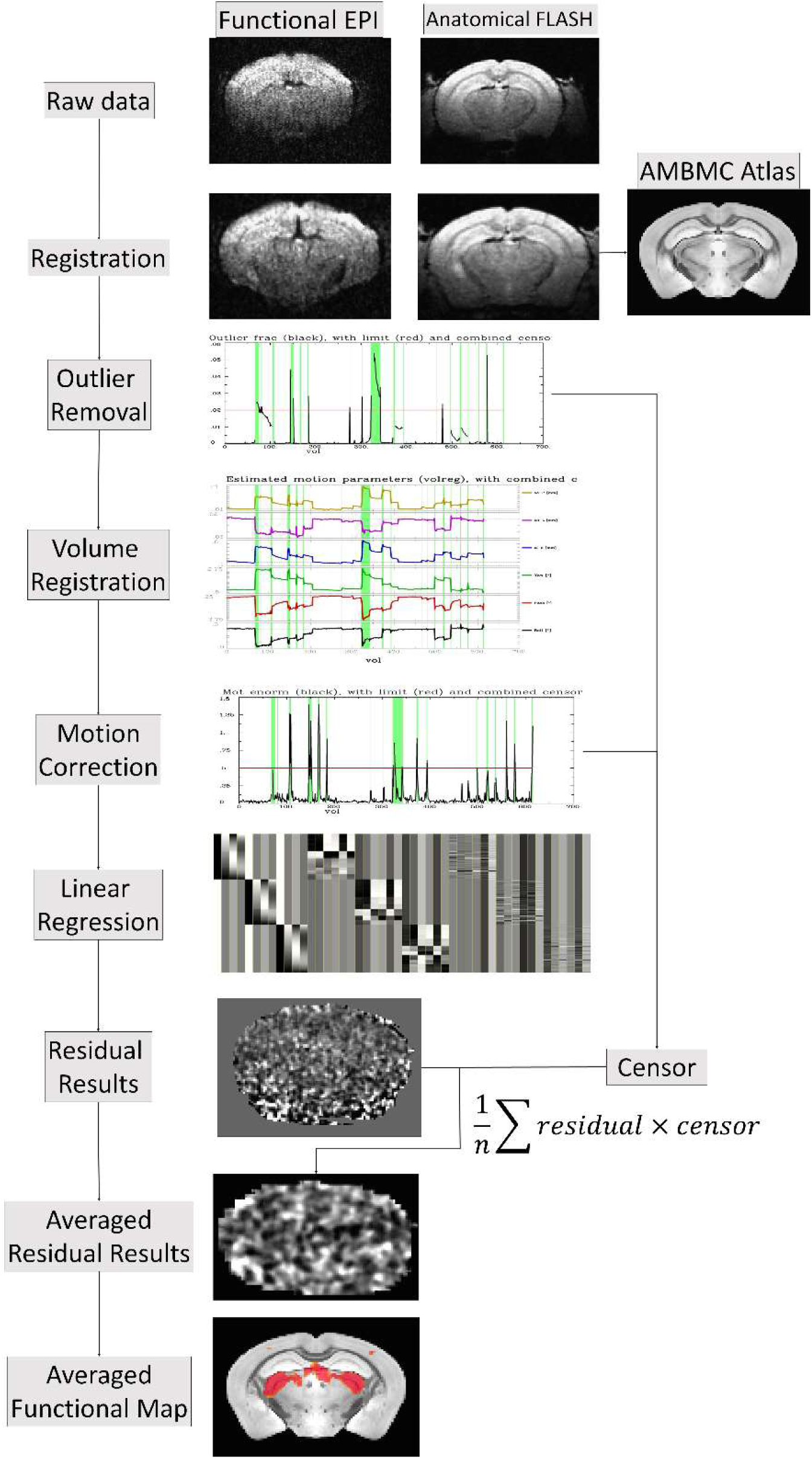
The processing pipeline of the awake mouse fMRI datasets. The workflow diagram is described through the following steps: raw data registration, outlier removal, volume registration-based motion file estimation, Motion removal, linear regression with the censoring function, functional map demonstration.

**Supp Fig 5.**
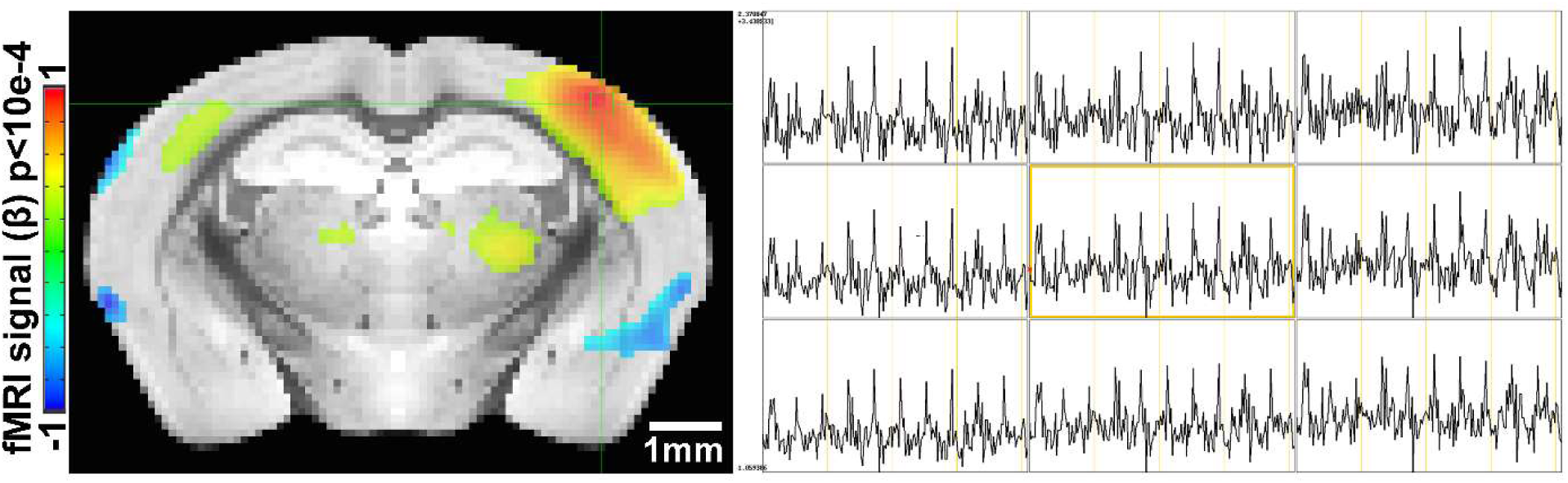
Time course from single experiment showing strong BOLD activations during stimulation paradigm. **Left**) BOLD activation map highlighting positive activation in the BC and selection of a 3x3 voxel display of time course data (green box). **Right**) Time course data taken from 3x3 voxel selection clearly showing 10 stimulations in each of the voxels shown.

**Supp Fig 6.**
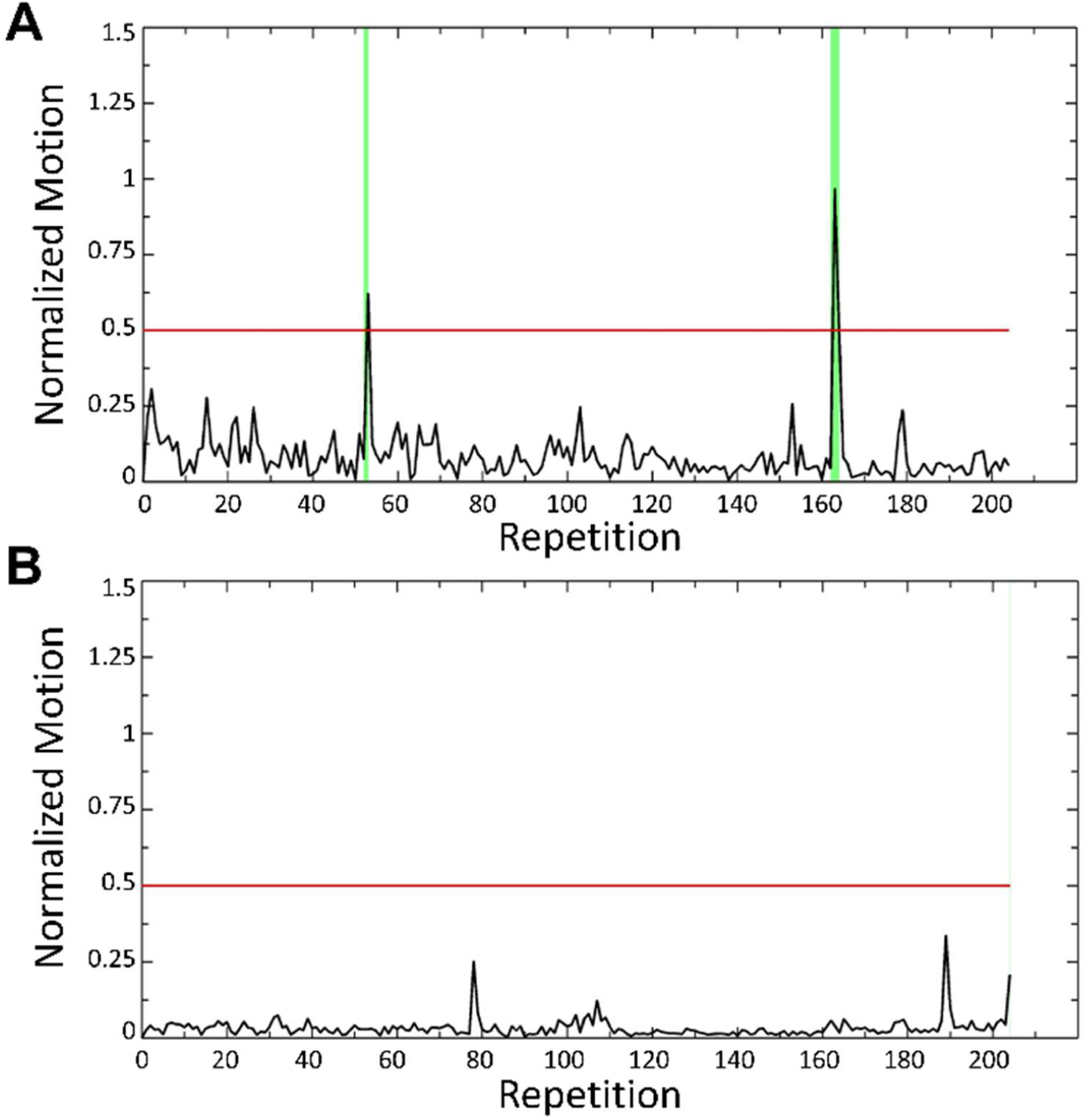
Figure showing motion data (.MOT file) of a representative mouse immediately following training (**A**) and after 1 month of scanning (**B**) indicating the amount of struggling the animal is doing during scanning. Struggling induced spikes are already substantially minimized following the training with only 5 volumes (2%) censored. After 1 month of scanning there was only one single repetition (repetition 204) at the end of the scan that was censored.

**Supp Fig 7.**
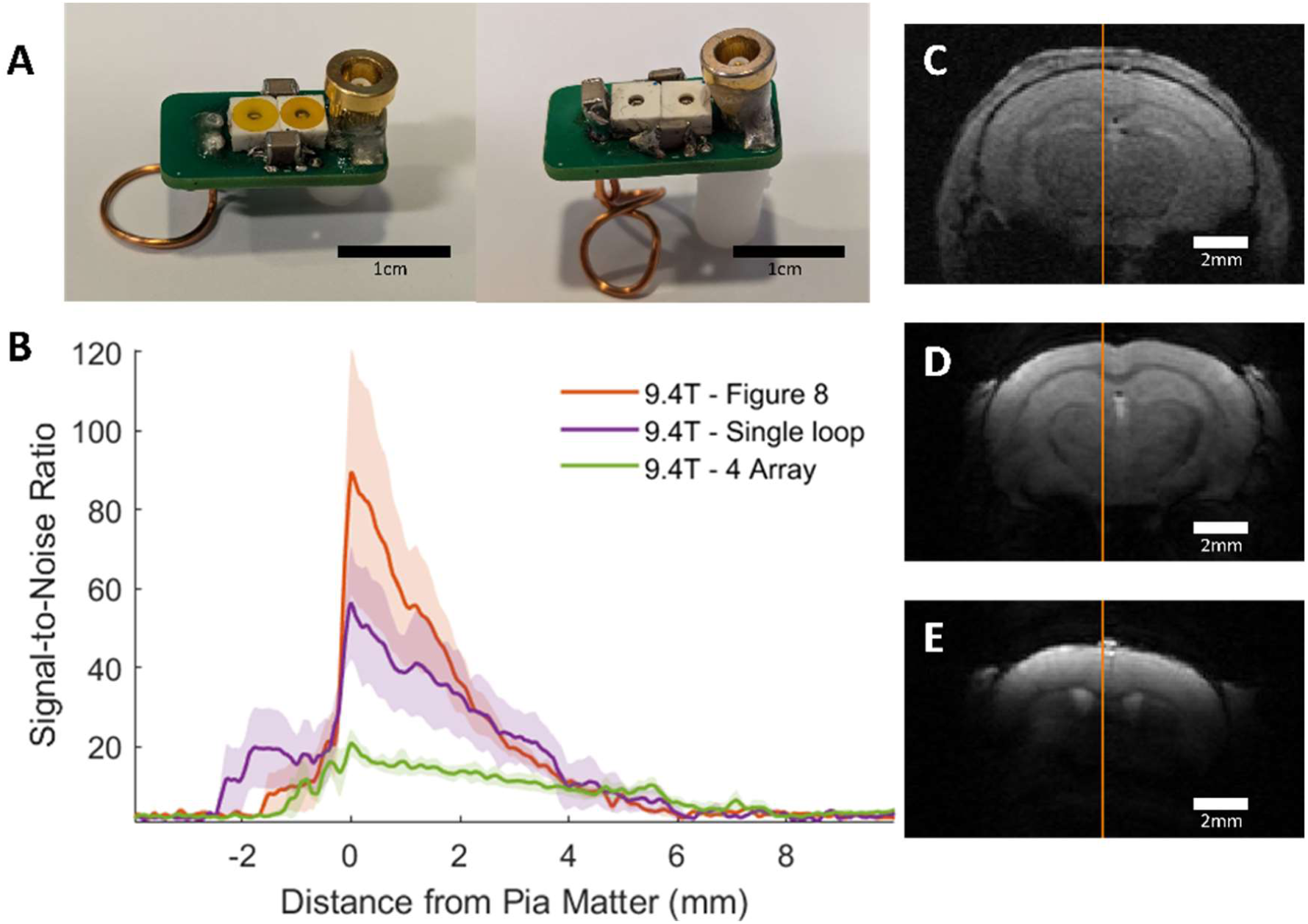
Comparison between implanted and commercial coils. **A** shows representative (unimplanted) coils in the single loop (**left**) and figure 8 styles (**right**). **Supplementary** Table 1 provides a parts list and cost for making these coils and **Supplementary** Figure 4 provides a circuit diagram to assemble. **B** presents the SNR line profile values as a function of distance from Pia Matter for each coil tested at 9.4T: commercial phased array surface coil (4 Array), implanted single loop, and implanted figure 8. SNR values were calculated by dividing the signal by the standard deviation of the noise. **C-E** shows a representative FLASH image with line profile of SNR measurements from each of the coils used to create the graph seen in **B**. Clear visual improvement in SNR can be seen in figures **C-E**. **C** – Commercial phased array. **D** – Single loop at 9.4T. **E** – Figure 8 at 9.4T. (N4 array = 6, Nsingle loop = 5, Nfigure 8 = 5)

**Supp Table 1.**
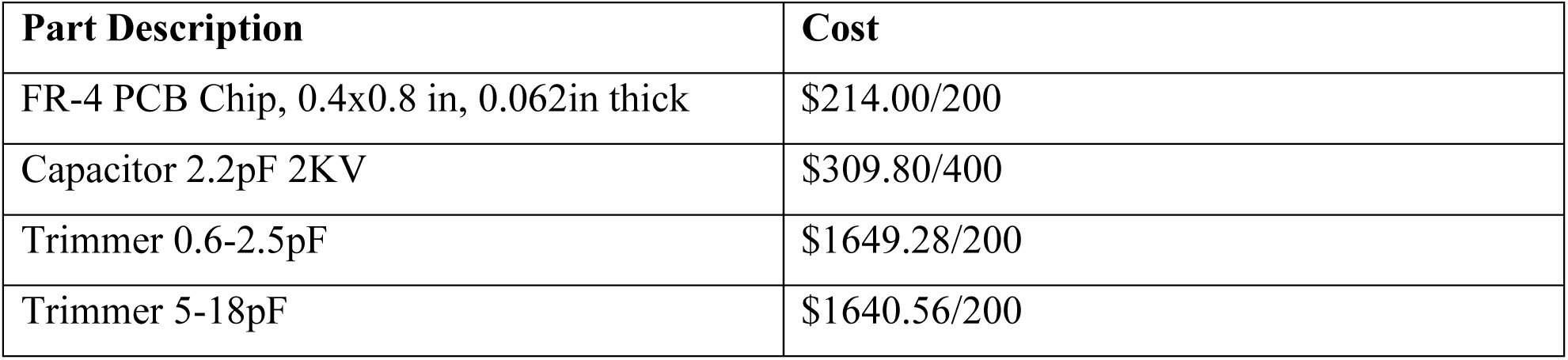
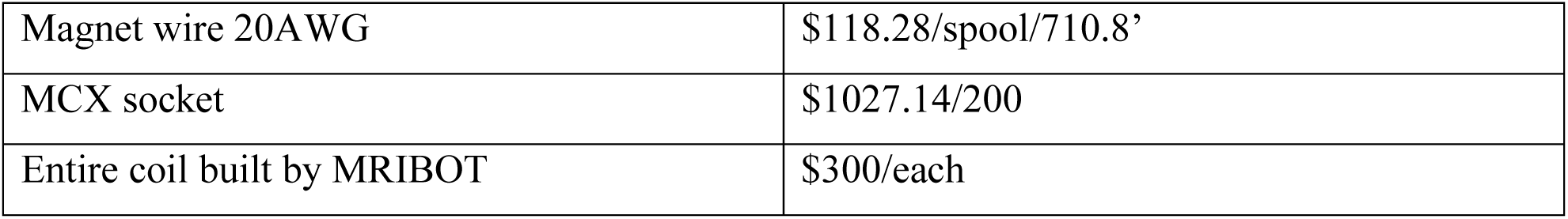
Parts list for construction of 200 ^1^H coils configured for 600MHz (14T).

## Supplementary Movie Legend

**Supp Movie 1.** The video shows how awake mouse is set up through the cradle using the implanted RF coil.

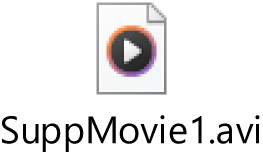

**Supp Movie 2.** The video shows the real-time EPI raw images from awake mice. The real-time tracer from the selected point demonstrates the time points with motion, as well as the motion-induced image distortion during awake mouse fMRI.

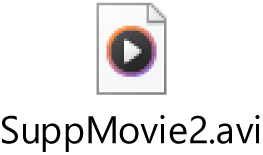

